# The Dementias Platform UK (DPUK) Data Portal

**DOI:** 10.1101/582155

**Authors:** S Bauermeister, C Orton, S Thompson, R A Barker, J R Bauermeister, Y Ben-Shlomo, C Brayne, D Burn, A Campbell, C Calvin, S Chandran, N Chaturvedi, G Chene, I P Chessell, A Corbett, D H J Davis, M Denis, C Dufouil, P Elliott, N Fox, D Hill, SM Hofer, M T Hu, C Jindra, F Kee, C H Kim, C Kim, M Kivimaki, I Koychev, R A Lawson, G J Linden, R A Lyons, C Mackay, P M Matthews, B McGuiness, L Middleton, C Moody, K Moore, D L Na, J T O’Brien, S Ourselin, S Paranjothy, K S Park, D J Porteous, M Richards, C W Ritchie, J D Rohrer, M N Rossor, J B Rowe, R Scahill, C Schnier, J M Schott, S W Seo, M South, A Steptoe, S J Tabrizi, A Tales, T Tillin, N J Timpson, A W Toga, PJ Visser, R Wade-Martins, T Wilkinson, J Williams, A Wong, J E Gallacher

**Affiliations:** Department of Psychiatry, University of Oxford; Swansea University Medical School, Swansea University; Cambridge University Department of Clinical Neurosciences and Cambridge University Hospitals NHS Foundation Trust, Cambridge; Population Health Sciences, University of Bristol; Department of Public Health, University of Cambridge; Faculty of Medical Sciences, Newcastle University; Department of Medical Genetics, University of Edinburgh; Centre for Clinical Brain Sciences, University of Edinburgh; MRC Unit for Lifelong Health and Ageing, UCL; Bordeaux Population Health, Université Bordeaux; Neuroscience, BioPharma R&D, AstraZeneca, Cambridge; College of Medicine and Health, University of Exeter; Oxford Academic Health Science Network, University of Oxford; MRC Centre for Environment and Health, School of Public Health, Imperial College London; Department of Neurodegenerative Disease, UCL Queen Square Institute of Neurology; King’s College London, London; Department of Psychology, University of Victoria; Nuffield Department of Clinical Medicine, University of Oxford; Centre for Public Health, Queen’s University Belfast; Department of Preventive Medicine, Yonsei University College of Medicine, Seoul; Institute of Epidemiology and Health, University College London; Translational and Clinical Research Institute, Newcastle University; Division of Brain Sciences and UK Dementia Research Institute, Imperial College; Medical Research Council; Department of Neurology, Samsung Medical Center, Sungkyunkwan University School of Medicine; Department of Psychiatry, University of Cambridge; School of Biomedical Engineering & Imaging Sciences, King’s College London; School of Medicine, Cardiff University; Institute of Health Science, Gyeongsang National University; Institute of Genetics and Molecular Medicine, University of Edinburgh; Usher Institute of Population Health Sciences and Informatics, University of Edinburgh; Department of Behavioural Science and Health, UCL; Centre for Innovative Ageing, Swansea University; UCL Institute for Cardiovascular Science; Laboratory of Neuro Imaging, USC; VU University Medical Centre, Maastricht University; Department of Physiology, Anatomy and Genetics, University of Oxford; Institute of Psychological Medicine and Clinical Neurosciences, and UK Dementia Research Institute, Cardiff University; Imperial College NIHR Biomedical Research Centre, Imperial College London; Health Data Research UK London at Imperial College London

## Abstract

The Dementias Platform UK (DPUK) Data Portal is a data repository facilitating access to data for 3 370 929 individuals in 42 cohorts. The Data Portal is an end-to-end data management solution providing a secure, fully auditable, remote access environment for the analysis of cohort data. All projects utilising the data are by default collaborations with the cohort research teams generating the data.

The Data Portal uses UK Secure eResearch Platform (UKSeRP) infrastructure to provide three core utilities: data discovery, access, and analysis. These are delivered using a 7 layered architecture comprising: data ingestion, data curation, platform interoperability, data discovery, access brokerage, data analysis and knowledge preservation. Automated, streamlined, and standardised procedures reduce the administrative burden for all stakeholders, particularly for requests involving multiple independent datasets, where a single request may be forwarded to multiple data controllers. Researchers are provided with their own secure ‘lab’ using VMware which is accessed using two factor authentication.

Over the last 2 years, 160 project proposals involving 579 individual cohort data access requests were received. These were received from 268 applicants spanning 72 institutions (56 academic, 13 commercial, 3 government) in 16 countries with 84 requests involving multiple cohorts. Project are varied including multi-modal, machine learning, and Mendelian randomisation analyses. Data access is usually free at point of use although a small number of cohorts require a data access fee.

## Introduction

Here we describe the Dementias Platform UK (DPUK) Data Portal (https://portal.dementiasplatform.uk/) (1). The Data Portal is a collaboration between DPUK and a growing number of cohort research teams who wish to make their data globally accessible. DPUK was established by the Medical Research Council (MRC) to accelerate the development of new treatments for dementia. The Data Portal is a component infrastructure of DPUK, designed to exploit the opportunities provided by cohort data to inform experimental medicine and to improve access to cohort data more widely. It is a data repository facilitating access to data for 3 370 929 individuals in 42 cohorts. The Data Portal provides a secure, fully auditable, remote access environment for the analysis of cohort data. The Data Portal supports the FAIR principles (Findability, Accessibility, Interoperability and Reusability) to improve the infrastructure supporting the re-use of data (2)

Arguments for multi-cohort-focused data repositories include: 1) as research questions focus on smaller effect sizes, access is required to data at-scale to achieve statistical purchase, 2) as emerging research questions become more complex, access to diverse multi-modal data is needed for rigorous hypothesis testing, 3) as scientific rigour increases there is growing recognition of the value of triangulation and replication using independent datasets, 4) as cohort datasets increase in size the transfer of large datasets is decreasingly feasible, 5) as cohort datasets become more complex the mastering of bespoke data models for survey, omics (genomics, proteomics and metabolomics), imaging and device data becomes burdensome, 6) as cohort datasets become more sensitive the non-auditable use of data is decreasingly acceptable. Whilst these issues can be addressed individually, the Data Portal provides an integrated solution.

The Data Portal is an end-to-end data management solution designed to support cohort data sharing. All projects utilising the data are by default collaborations with the cohort research teams generating the data. Although it has the three core utilities of data discovery, access, and analysis, to achieve these it operates across seven layers (Figure).

### Layer 1- Data Ingestion

The data journey begins with upload to the Data Portal. Datasets and data dictionaries are received from cohorts on an ‘as-is’ basis along with other supporting documentation. The Data Portal operates within the UK Secure eResearch Platform (UKSeRP) environment according to ISO 27001 (3) and operates exclusively as a data processor according to the UK Data Protection Act 2018 (4) and EU General Data Protection Regulation 2016 (5). DPUK facilitates the legal engagements necessary for data transfer into the Data Portal on behalf of data controllers, by ensuring robust contractual arrangements are in place as an overarching mechanism for data governance and use (6). In practice the data controller is considered to be the principal investigator for the dataset. A dataset is removed from the Data Portal upon receipt of a wet-ink signature request from the data controller.

### Layer 2- Data curation

Upon receipt, ‘native’ data are curated to a common data model (C-Surv). The C-Surv ontology is designed to simplify the analytic challenge of working across multiple datasets and multiple modalities by providing standard structure, variable naming and value labelling conventions. Other data models, CDISC (7), OMOP (8), or HPO (9) involve structural complexity that is rarely relevant to cohort-based analyses. An example variable name using the C-Surv model is given below:

GEN05_PAINCHESTEVR_0_1

The cohort is identified using a three digit alphabetic character (GEN for Generation Scotland), and category-level data by a two digit numeric character (05 for physical health status). The measure is described by an alphanumeric acronym (PAINCHESTEVR for: Do you ever get pain or discomfort in your chest?). This is followed by an integer giving the number of repeat measurements within a study wave (_0 indicates there were no repeat measurements). Finally, an integer suffix indicates study wave (_1 for recruitment, _2 for the first follow-up, etc.). This data model is a contribution to a wider debate on how global access to cohort data can be achieved.

C-Surv is optimised for the analysis of ‘flat-file’ data. Higher order data must be pre-processed prior to curation. The XNAT (10) imaging platform is used to receive and process DICOM and NIfTI files. For genetics, variant call format and allele frequency data may be uploaded. C-Surv is also designed to improve the efficiency of data selection and management from the applicants and administrators’ perspectives. There are 22 categories and 132 sub-categories, which may be used for data selection as an alternative to individual variable selection. The ontology is machine-readable for mapping to other ontologies. Researchers may request access to either native or curated data. Standardisation does not imply harmonisation i.e. comparability (equivalence) of values and/or distributions across variables. Data harmonisation is implicitly purpose specific. DPUK undertakes a very limited harmonisation programme solely to support data discovery (see Layer 4). Data curation is resource intensive and is ongoing. To accelerate the process we are developing machine-learning approaches, which currently achieve an 80% accuracy.

### Layer 3- Interoperability

We anticipate a global mixed-model data access environment that respects sovereign boundaries and requires the highest levels of security. In practice, this involves the integration of individual participant data across datasets (‘pooled’ analyses) and the integration of summary data across datasets (‘federated’ analyses). Both models require interoperability between data platforms across national boundaries. DPUK supports the development of interoperability in several ways. In partnership with UKSeRP, we provide access to our software solutions to other data platforms to facilitate architectural compatibility. We also engage with other data platforms to develop interoperability across architectures. In collaboration with Gates Ventures and other data platforms, we are working to develop a high-level data-gateway for cross-platform data access.

### Layer 4- Data discovery

A tiered data discovery pathway begins with the Cohort Matrix (11) which provides a high-level comparison of data availability for each cohort. The Cohort Directory (12) enables detailed exploration across cohorts using a range of metadata categories. The Cohort Explorer (13) provides access to a set of 30 variables harmonised across cohorts to enable feasibility analysis. Adding cohorts to the Cohort Explorer is ongoing. As the Cohort Explorer provides access to harmonised data, login to the Data Portal is required. Data discovery is a dynamic area and the development of increasingly ergonomic and intuitive tools is anticipated. This is likely to include the identification of high priority variable sets and variable selection algorithms through the analysis of user-activity and the development of application programming interfaces (APIs) to rapidly reproduce data selection (see Layer 7). DPUK welcomes user feedback and collaboration to inform an ongoing programme of discovery tool development.

### Layer 5- Data Access brokerage

Automated third-party brokerage, allowing submission of a single data access request to multiple data controllers, reduces the researcher administrative burden. Standard cross-cohort access agreements and streamlined decision-making procedures simplifies the application and approvals process for all stakeholders. Access requests can be specified at the level of detail required by the data controller and if approval is granted, a standard access agreement must be completed by both parties prior to access being granted. The application form is a synthesis of key issues that are addressed by most, if not all, the individual cohort access management procedures, fail-fast criteria have been developed to facilitate rapid and transparent decision-making by data controllers. These comprise the proposal not being in the public interest, potential identifiability of participants, no clear scientific rationale, no appropriate analysis plan, or a conflict of scientific interest. Upon access approval by a cohort data access committee (DAC), completion by the applicant of a data access agreement is required prior to access being granted. Upon receipt of a completed data access agreement, data access is granted within several days for any one cohort (14). Data access is normally free at the point of use with a small number of cohorts levying a data access fee (Table).

### Layer 6- Data analysis

The analysis environment is VMware based with each researcher allocated their own private personal ‘lab’ (desktop) into which approved data are moved, and from which they can access a range of pre-loaded generic and specialist software packages. Bespoke software may also be uploaded upon approval. VMware clients are available for a range of operating system including Windows, Linux, IOS, Mac, and Android. A standard desktop is provided for researchers seeking to access standard phenotypic data on the Portal. The standard desktop has the following specification: Windows 7/10, 8GB RAM, and 4 CPUs. It is pre-loaded with R, RStudio, SPSS, SAS, Stata, Python, Eclipse, MATLAB, SQL Server Management Studio, Microsoft Office. Statistical software, such as R and Python, can connect to its official library/package/index directory to enable configuration of software on a per-user basis. Larger desktops are available on request (32GB RAM with 8 CPUs, 128GB RAM and 16CPUs) which are more suitable for large-scale multi-modal analyses such as omics and imaging analyses, and for machine learning.

Storage is scalable according to the study requirements, with basic access to data stored on our systems free to all users. Studies needing unusually large amounts of storage to ingress their own at-scale data will be subject to potential charges which will be discussed with researchers as part of the desktop set-up. High Performance Computing and non-Windows operating systems are available upon request. The Data Portal also offers consortia-based workspace; providing an independent and transparent storage, access and analysis solution for use by multiple institutions.

Two-factor authentication is required to access approved datasets. This involves the provision of a username with password creation, and an authentication code generated by an app on a mobile device of the applicant’s choosing. Data may not be removed from the Portal. However, tables, graphs and scripts for export are submitted to the data export panel for approval. Manuscripts may be prepared in the Data Portal so that collaborators who are registered users may contribute without the need for manuscript download. A facility for import is also available, enabling researchers to upload scripts and additional datasets from outside the Data Portal to reside within their approved DPUK datasets.

### Layer 7- Knowledge environment

Cohort data grow in quantity and can be complex, but ultimately there is the need for the transition from data to knowledge and insight. In response to this challenge, the Data Portal is developing a Knowledge Hub to organise and make information readily accessible on the key activities of the Data Portal. The immediate purpose of the knowledge Hub is to enable collaborative knowledge building; enabling researchers to understand what is available, what is known, and where there are knowledge gaps. It also serves as a knowledge preservation repository enabling the storage of linked datasets and analysis code with persistent identifiers for rapid access and replication. DPUK welcomes collaboration in the development of this hub.

Application for access can be made through the Data Portal https://portal.dementiasplatform.uk/ (15)

## Results

### Data resources

DPUK collaborates with 42 clinical and population cohorts (Table). Of these, 24 (n=1 410 302) have either completed or are in the process of uploading full or partial datasets and 18 (n=1 960, 627) upload data on a per-project basis. The data are diverse. Clinical cohort studies include familial disease cohorts (16, 17), disease focused patient cohorts (18, 19), ageing focused population cohorts (20-23), re-purposed cardiovascular cohorts (24, 25), birth cohorts (26, 27), repurposed cancer cohorts (28, 29), and disease agnostic cohorts (30-32). In terms of real world evidence, the Data Portal provides access to an e-cohort (SAIL-DeC) covering health records for 1.2m individuals including 130k dementia cases (33). Data availability varies according to cohort but includes epidemiologic survey, imaging, genetics, and linked administrative data. The Data Portal provides a link to the UK-CRIS Network for natural language processing of 2.5m UK mental health records (34). Overall, these cohorts and the data modalities represented, provide an unusually rich and complex data environment.

### Data resource use

From launch in November 2017 to end of January 2019, 160 project proposals involving 579 individual cohort data access requests were received. These were received from 268 applicants spanning 72 institutions (56 academic, 13 commercial, 3 government) in 16 countries with 84 requests involving multiple cohorts.

Of the 160 project proposals, 40 failed triage due to under-specification and were not forwarded to cohort DACs. The 120 projects that were forwarded to DACs involved 396 individual cohort access requests, 19 of which were withdrawn by the applicant pre-approval. Of the 377 access requests considered 171 (45%) were approved, 80 (21%) were declined, 78 (21%) were closed following non-response after 200 days from either the applicant or the cohort, 19 (5%) ceased after approval and 29 (8%) are in process. Of the 80 declined requests, 49 (61%) were due to the data requested either not existing or not being relevant for the analysis.

The data requests were diverse. Projects include Mendelian randomisation, imaging, psychometric, and machine learning studies, alongside risk stratification studies and dementia prediction and diagnosis modelling. Other studies are less dementia specific including trajectories of longitudinal assessment of comorbidities between mental health, hormonal indicators and cognitive change; the impact of child adversity on adult outcomes; the longitudinal tracking and determinants of well-being and cognitive performance; and successful cognitive ageing in 90+ year olds.

To facilitate innovative and exploratory analyses, the Data Portal can be configured as a ‘sandbox’ environment, without compromising data security or cohort governance principles. An example of this was the hosting of a three-day multidisciplinary workshop (datathon) for 40 data scientists at the Alan Turing Institute utilising the Deep and Frequent Phenotyping pilot study data (35). These multi-modal data include Magneto-encephalography (MEG), Positron Emission Tomography (PET), structural and functional MRI, ophthalmology, gait, and serial cognitive and clinical assessment. The Data Portal was used to host 25 data scientists in a datathon at the University of Exeter and Swansea University, during which traditional regression and machine learning procedures were used to interrogate the data within the virtual desktop interface. A datathon at the University of East Anglia hosted 60 attendees, mostly early career researchers, using data from the Deep and Frequent Phenotyping Study (35), the English Longitudinal study of Ageing (20), and the Caerphilly Prospective Study (36). The datathon programme is ongoing with a further 6 planned for 2019. The sandbox environment is also suitable for teaching and training purposes as it enables access to standardised datasets by multiple users within a secure environment.

### Portal performance

Data Portal utility varies according to the extent that standardised procedures are used. Currently, 38 cohorts (90%) accept our standard legal agreements, 28 cohorts (67%) accept our streamlined data access process, with a further 13 accepting a DPUK application as an initial enquiry. For variable selection, 22 (52%) cohorts accept the use of our variable selection tools with the remainder requiring the use of cohort-specific variable selection tools (Table).

Triage is rapid, usually upon receipt. Cohort DAC decision times vary according to the complexity of the project, clarity of the proposal, and the DAC meeting frequency. DPUK policy is to consider a project closed (in effect expired) if there has been no response from the cohort DAC within 200 days. Our target DAC response time is 28 days. Current performance has a median value of 28 days (mean=40). Time from approval to data access is dependent on the completion of the data access agreement by the applicant. Where data are already uploaded to the Data Portal access is typically given with 24 hours of receiving the completed Data Access Agreement. Where data are uploaded on a per-project basis, access is dependent on the cohort data management process. Typically, upload of native data (not curated by DPUK) occurs within 2 weeks and can be made accessible to the researcher immediately.

## Discussion

Rapid data access continues to be a challenge for researchers, funders, and governments. The DPUK Data Portal is one of several initiatives to address this issue. In the dementia space, other data platforms are available. The JPND Global Cohort Directory (37) provides contact details for 175 cohorts (n=3 586 109) whilst the IALSA Network (38) provides details for 110 cohorts (n=1 485 410). More sophisticated and convenient data discovery tools are provided by GAAIN (39) with 47 cohorts (n=480 020). GAAIN also offers centralised processing for selected datasets. EMIF-AD (38) offers a comprehensive data harmonisation programme for a selection of their 60 catalogued cohorts (n=135 959) and 18 electronic health records datasets (n=65M). For selected datasets EMIF-AD provides centralised cohort data processing facilities through the TranSMART platform (40).

### Data Portal Benefits

However, in providing independent trusted third-party data management, the data repository approach has several benefits. Although the Data Portal is optimised for dementia, its remote access model provides a generic solution that can be adapted to any outcome for which relevant data are available. Specifically, by automating, streamlining and standardising procedures, decision-making can be more precise, quicker, and more transparent, and the practice of repeated data transfers avoided.

For cohort research teams a repository substantially reduces the administrative burden of routine data management tasks, particularly avoiding repeated data transfers. This assists cohorts in meeting their data sharing obligations, and releases cohort-based data scientists to focus on increasing the scientific value of their data.

For researchers a repository provides a single point of contact and a single standard procedure for accessing multiple independent datasets. Within the Data Portal this supports data discovery, feasibility modelling, access management and analysis. The use of a common data model reduces data preparation times and reduces uncertainty due to unnecessary variation in data pre-processing.

For the research community as a whole, a repository facilitates knowledge management including the linking of code-to-dataset through the use of persistent identifiers (DOIs), the development and discoverability of APIs for rapid replication, and the improvement of the end-to-end user experience through adaptive learning.

Repositories have an important role in the democratisation of science. The Data Portal allows remote and secure data analysis, enabling the best ideas to be tested in high quality data from any location with suitable connectivity. This is fundamentally empowering to researchers without access to high-end computing, particularly in developing countries.

### Data Portal Challenges

New infrastructure requires new working practice, which brings challenges to platform operators, data controllers and data users.

Operational challenges for platform operators focus on improving the user experience and increasing the scientific opportunity. Data platforms are complex infrastructures that integrate the highest levels of security and governance requirements for accessing large, complex, and non-standard datasets. They are also integrated within larger computing facilities which require specific resource allocation for computer intensive operations.

In this context, improving the user experience is an incremental process. For example, improving data discovery and visualisation tools to improve the precision of data access requests, streamlining decision making to reducing approval times, and developing links to similar resources globally. All involve engagement with different stakeholders, each of which have different expectations and requirements.

For some data controllers the repository ‘trusted third-party’ approach to data management involves uncertainty. Whilst the practice of managing access through repeated data transfer may be labour intensive, it is familiar. Uncertainty surrounds the legal status of EHR data accessed via the platform, quality of the proposed science, the relevance (and existence) of the requested data, and potential conflicts of scientific interest. The legal status of accessing EHR data linked to research data using third party platforms is a grey area in the UK. Although there is no change in data controller, and no forward sharing to a third-party data controller, current UK data sharing licenses have not anticipated the Data Portal solution. Discussions are underway in the UK data community to resolve this issue. Wider access also brings reduced control, and data controllers can be concerned over the reputation of their datasets if used for egregious science. In response DPUK strongly advises a collaboration with cohort research groups as this is the quickest way to understand a dataset and receive expert advice on how to approach its analysis. To facilitate good science it is not unreasonable for cohort research teams to insist on a collaborative approach. That 61% of declined access requests (see Data Resource Use) were due to the data requested either not existing or not being relevant for the analysis is to an extent a matter of due diligence on the part of the applicant, but it can also be a matter of judgement. The impact of conflict of interest depends largely on the perspective of the DAC. In an age of increasingly large and complex analyses, many cohorts are underpowered for emerging research questions. Cohort research team interests can be well-served though transparent collaborative relationships. To reassure data controllers, their approval is required for all data access, all use is auditable, data cannot be downloaded, and data are erased upon request.

For researchers, accessing data via a repository may be less convenient than through their host institution. However, the additional security measures are not onerous and are balanced by improved data access. A secure repository requires an account and user authentication. Whilst these procedures may be simple (two-factor authentication takes ≈10 seconds), they require a level of familiarity before becoming second nature. For access approval, epidemiologic, imaging, and genetic communities each have different conventions over what constitutes an appropriate data request. By streamlining and standardising the application process DPUK encourages coalescence around the development of common standards that are widely accepted. The informatics infrastructure required by these groups also varies in terms of operating systems, compute power, and software. Our response is to provide flexibility. Standard environments are available that suit the requirements of most users, but bespoke environments are built on request including access to high performance computing.

In the pursuit of scientific rigour, increased data access is only part of the equation. Also required is a commensurate development in analytic competence. That 40 project proposals failed a not very demanding triage process, suggests that some analysts are unaware of the value and complexity of the data resources that are available. Although DPUK can increase the opportunity for training via the Data Portal, there is a wider responsibility to up-skill analysts to take advantage of this.

### Looking ahead

The mission of the Data Portal is to increase the realised scientific value of cohort data. Future developments include developing linkage to electronic health records. In partnership with Health Data Research UK, DPUK is developing a centralised system enabling DPUK cohorts to link research data to routinely collected data. In partnership with Gates Ventures, DPUK along with other platforms is developing global connectivity. DPUK is also working with academic and industry partners to provide digital phenotyping using devices such as smart phones, for collaborating cohorts. A third development is the use of cohort data to risk-stratify for trials recruitment. For dementia, there is a growing need for risk stratification by phenotype according to genotype in order to recruit to highly targeted experimental medicine studies. Cohorts, by virtue of lead-in data are well positioned to support these studies. DPUK collaborates with the Airwave and Health-Wise Wales cohorts to provide genotyping and cognitive testing for experimental medicine recruitment. DPUK welcomes extending this collaboration to other cohorts.

### Conclusions

The DPUK Data Portal was established by MRC to support the development of new treatments for dementia by using cohort data to inform experimental medicine. It is recognition of the unique value of cohort data and a contribution to the wider debate on how best to support cohort studies and facilitate their contribution to the wider research environment. By streamlining procedures for cohort research teams, increasing data accessibility for researchers, reducing costs, and adding value, the Data Portal is an investment in the future of cohort studies.

Applications for access can be made through the Data Portal https://portal.dementiasplatform.uk/

## Acknowledgements

DPUK would like to express gratitude to:

Cohort members and their research teams for generously making data available

EMIF-AD for providing access to their data catalogue and supporting software

Professor Ian Deary, Dr Declan Jones, and Professor Sir Simon Lovestone for their contribution to this paper from their support in the DPUK Executive Team.

## Declarations

### Availability of data and material

Not applicable

### Code availability

Not applicable

### Authors’ contributions

All authors contributed to the conception, creation and development of all the themes of the DPUK itself, including this Data Portal. Material preparation, by John Gallacher, Sarah Bauermeister, Chris Orton, Simon Thompson. The first draft of the manuscript was written by John Gallacher and all authors commented on previous versions of the manuscript. All authors read and approved the final manuscript.

### Ethics approval

Not applicable

### Consent to participate

Not applicable

### Consent for publication

Not applicable

### Availability of data and material

Not applicable

### Code availability

Not applicable

---

**Table:**
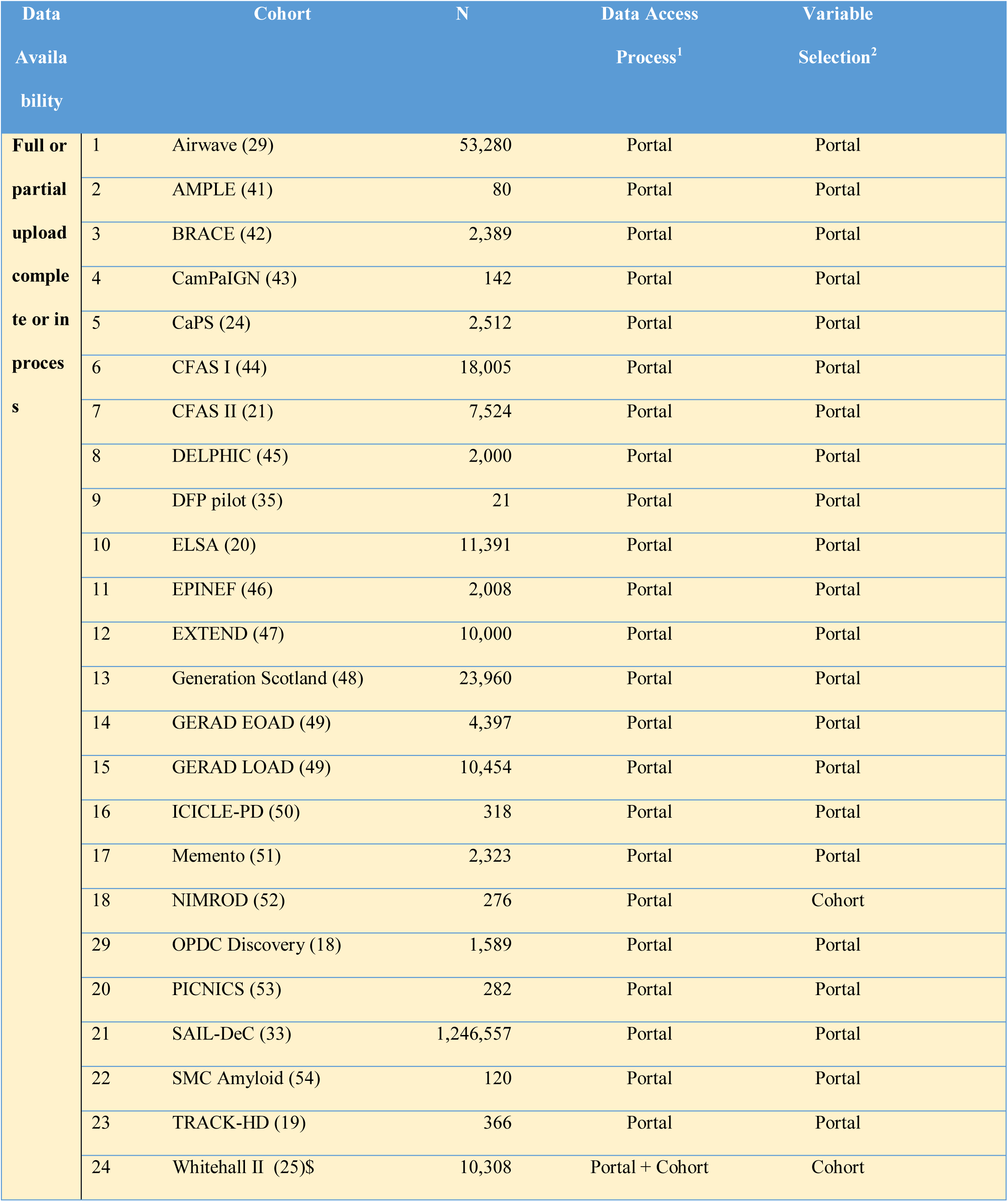
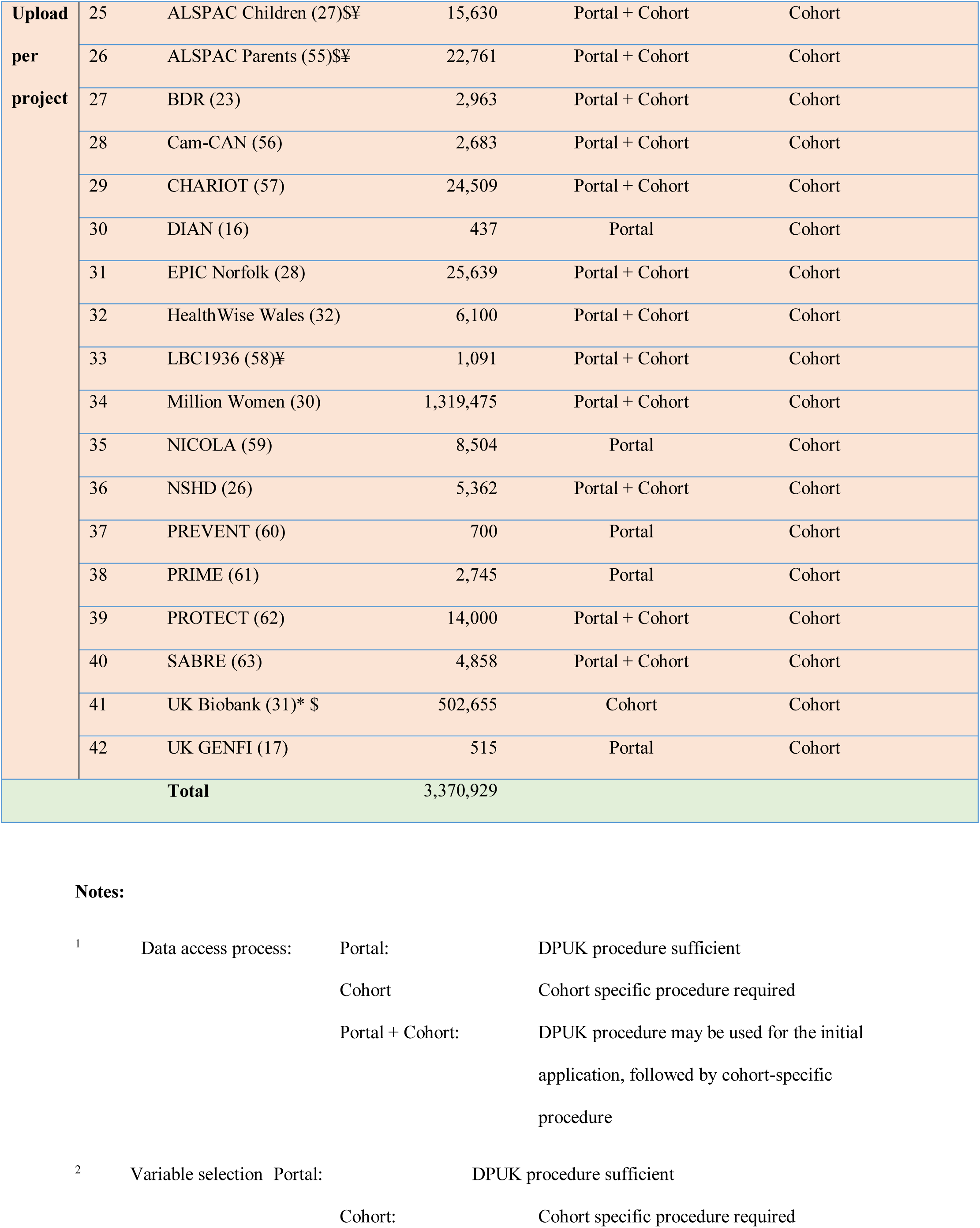
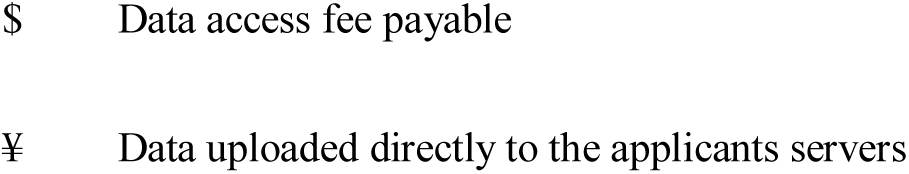
Collaborating cohorts.

**Figure:**
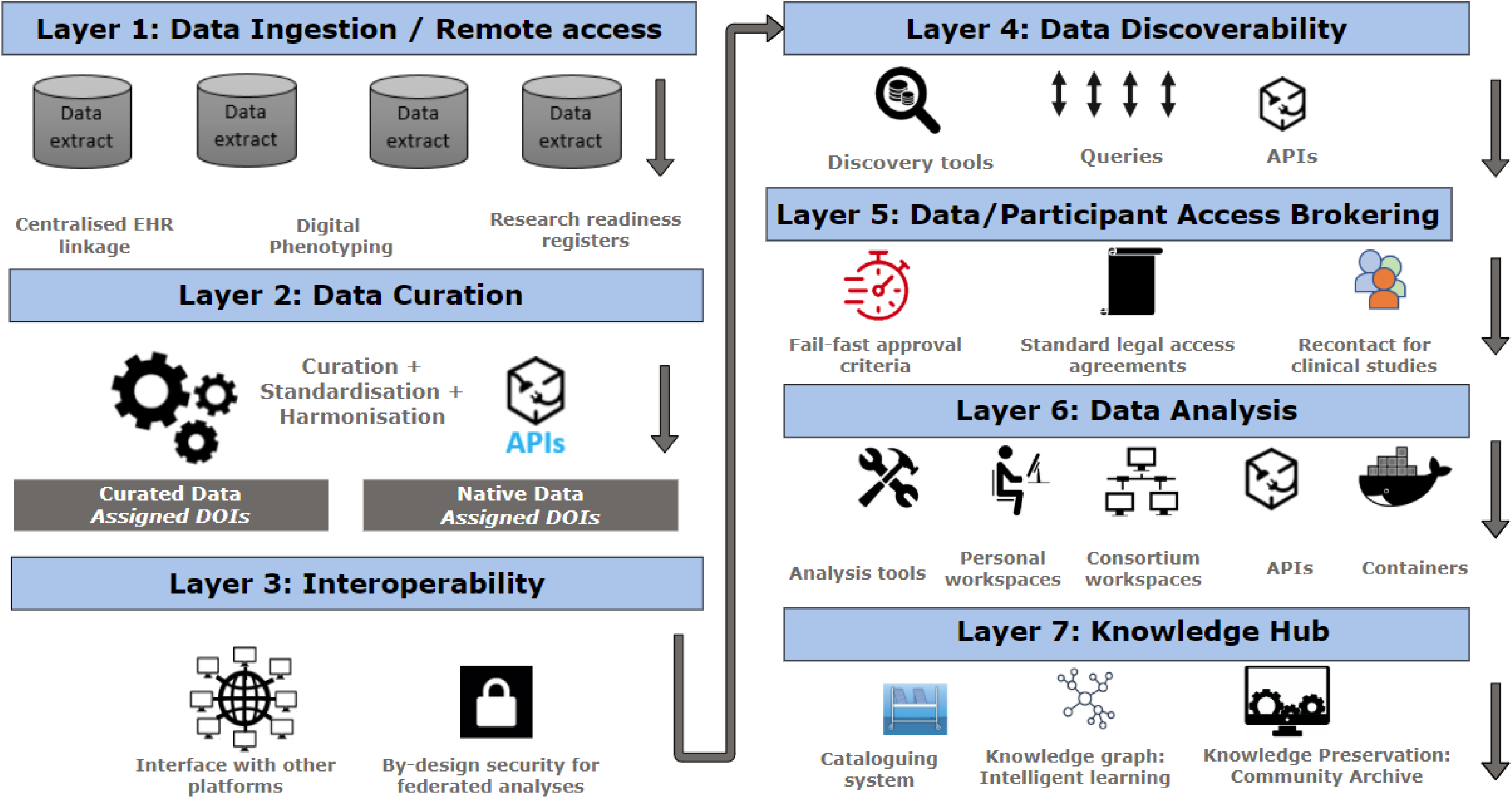
Data Portal architecture.

